# Microfluidic Formation of Ultrathin, Handleable Collagen Sheets Exhibiting Toe-heel Tensile Behavior

**DOI:** 10.1101/2024.09.27.615530

**Authors:** Yuming Zhang, Shashi Malladi, Bangan Wang, Elliot L. Chaikof, Axel Günther

## Abstract

The extracellular matrix (ECM) of cardiovascular tissues displays a non-linear, strain-dependent elastic modulus, attributed to the hierarchical organization of collagen. At low loads, these tissues exhibit compliance, permitting contraction or dilation, while at high loads, they stiffen considerably, increasing their mechanical strength by at least tenfold. Although collagen gels are widely used in 3D cell culture, tissue engineering, and biofabrication, current engineering techniques fail to replicate this hierarchical organization at the microscale. As a result, they lack both the non-linear tensile behavior and the physiologically relevant strength of native tissues. To address this limitation, we present templated collagen sheets that are 1.8 microns thin and 10 mm wide that demonstrate non-linear tensile behavior. Collagen sheets are obtained from an acidic collagen solution via a microfluidic flow focusing process, incorporating and subsequently removing emulsified oil droplets (mean diameters 2.1 microns and 5.0 microns, volume concentration 2.25%). Templated collagen sheets exhibit a two-fold increase in fibril alignment dispersion compared with non-templated ones. When assessed along their length, the Young’s modulus of templated sheets increases 62-fold at 90% failure strain, closely matching the properties of native load-bearing tissues. We anticipate that these ultrathin templated collagen sheets will have broad applications as a substrate material for the bottom-up fabrication of load-bearing biomaterials and tissue structures for in vitro applications and implantation.

## 1. Introduction

A distinguishing feature of many cardiovascular tissues, such as large blood vessels, is their strain-dependent elastic modulus, illustrated by a J-shaped curve of tensile stress as a function of applied strain (**Fig. 1A**).^[1-3]^ This non-linear relationship is characterized by a shallow slope at small strain values, typically below 10% of the strain to failure (STF), referred to as the “toe” region, and a steep slope at high strain values, typically above 90% failure strain, known as the “heel” region. The strain-dependence of the slope, or Young’s modulus (*E*), can be attributed to the unique spatial organization of load-bearing extracellular matrix (ECM) constituents, including fibrillar-forming type I collagen and elastin.

**Figure 1.**
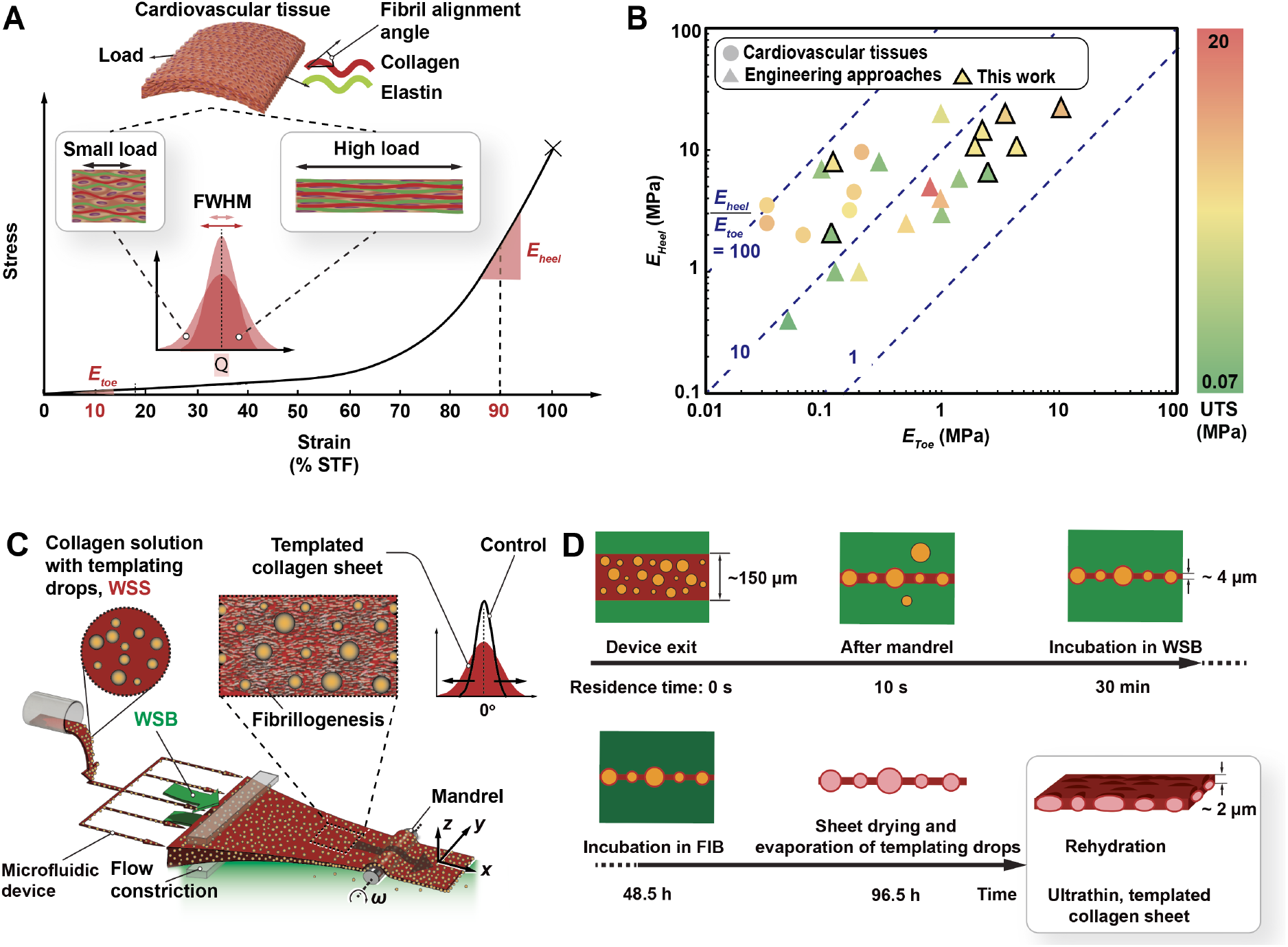
Multiscale organization of collagen in load-bearing soft tissues and engineered biomaterials. (A) Schematic illustration of multiscale fibrillar organization in arterial tissues at small and high loads. (B) Strain-dependent Young’s moduli in cardiovascular tissues and engineered biomaterials (for detailed literature references see **Section S1**, Supporting Information). (C) Schematic illustration of presented wet-spinning approach for preparing aligned, templated collagen sheets. (D) Post-processing protocol for preparing ultrathin, templated collagen sheets.

In the toe region, collagen fibers are undulating and exhibit greater fibril alignment dispersion, resulting in a small Young’s modulus, *E*_*toe*_.^[4, 5]^ In this low-strain configuration, fibrils straighten with minimal resistance to applied load. As the strain increases, more fibrils straighten and align, becoming load-bearing.^[6-8]^ In the heel region, close to the ultimate tensile strength (UTS) at the STF, the Young’s modulus, *E*_*heel*_, is one to two magnitudes higher than *E*_*toe*_, with most fibrils straightened and minimal fibril alignment dispersion.

Native cardiovascular tissues exhibited complex mechanical properties. During load bearing, *E*_*toe*_ is 0.033 - 0.2 MPa, while *E*_*heel*_ is >1 MPa, resulting in ratios *E*_*heel*_/*E*_*toe*_ from 30 to 90.^[9]^ The UTS varies significantly across different tissues: 0.4 - 3 MPa in veins,^[10, 11]^ 1.5 - 4 MPa in arteries,^[6, 12]^ and 8 - 37 MPa in human heart valves (**Figs. 1B**).^[9, 13-16]^ Additionally, the STF of such tissues ranges from 20% in veins and exceeds 240% in arteries.^[17]^

Various techniques, including micromolding,^[18, 19]^ strain conditioning,^[20]^ and chemical/thermal conditioning,^[21, 22]^ have been explored to replicate the non-linear tensile behavior of collagen that is observed in intact tissues. In addition, bottom-up biofabrication methods have been used to produce collagen-based fibers,^[19, 23, 24]^ sheets,^[25, 26]^ and tubular structure.^[27-34]^ A common goal of many studies is to emulate the collagen fibril dispersion of native tissues. However, previously obtained structures either exhibit high compliance while lacking tensile strength, or display high stiffness irrespective of the applied strain, and therefore offer poor compliance (**Fig. 1B**). To truly mimic the structural behavior of cardiovascular tissues, a hierarchically organized biomaterial is needed that due to its strain dependent elastic modulus recapitulates *both* the high compliance and strength of intact cardiovascular tissues.

A class of bulk materials that is known to provide non-linear tensile properties are cellular solids that incorporate voids into a solid substrate, often a synthetic polymer.^[35]^ At low strain values, where the internal pores begin to collapse, cellular solids exhibit small *E*_*toe*_ values, due to the buckling of the wall structures in between pores. As strain increases and with it the collapse of the pores, the internal wall structures align with the direction of strain, leading to higher *E*_*heel*_ values.^[36]^ While this concept holds significant promise, it has to our knowledge not yet been applied to spatially constrained biomaterials.

Here, we report the biofabrication of ultrathin collagen sheets with high fibrillar alignment and density via the combined effect of hydrodynamic confinement, macromolecular crowding and strain. A multiscale fibrillar organization with increased fibrillar dispersion angle is induced by the void regions left by templating oil droplets after removal. The obtained collagen sheets exhibit non-linear stress strain behavior and strength similar to the ones of intact cardiovascular tissues.

## 2. Results and Discussion

### 2.1. Preparation of Templated Sheets

A wet spinning solution (WSS) was prepared that consisted of 5 mg ml^-1^ acidic rat tail type I collagen (for details on collagen extraction and WSS preparation, see **Section S2-3**, Supporting Information) and oil droplets (concentration 2.3 v/v%, mean diameters 2.1 µm or 5.0 µm, see **Section S4** for details of templating emulsion preparation). The surfactant, polysorbate 80, added at 3.0 v/v% into the aqueous phase and stabilized the emulsion produced during microfluidic droplet generation (critical micellular concentration = 0.012 mM).^[37-39]^ The silicone oil droplets are neutrally buoyant (density 0.91 g/cm^3^, viscosity 5.0 × 10^−6^ m^2^s^-1^ at 25 °C) and chemically inert. The WSS and the wet spinning buffer (WSB) were supplied to separate inlets of a three-layered microfluidic device as described by Malladi *et al*. (**Fig. 1C**).^[25]^. At the device exit, a uniform emulsion-containing collagen sheath emerged and was hydrodynamically focused between two WSB streams while passing through an attached flow constriction. The emerging sheet floated at the WSB surface while undergoing pH-induced gelation, as well as osmotic removal of water. The sheet was strained by a mandrel that rotated with angular velocity *ω*, located 55 mm downstream the exit of the 7.6 mm long flow constriction. ^[25]^ To characterize the magnitude of straining we introduce a dimensionless parameter

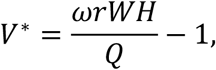

where *ω* · *r* is the linear velocity at the mandrel surface, *W* · *H* is the cross-sectional area of the flow constriction, and *Q* is the total volumetric flow rate of WSS and WSB combined. The strain applied by the mandrel resulted in a significant reduction in the sheet width up to 60%, compared to the width of the microfluidic device exit.

Based on our previous work on non-templated sheets,^[25]^ we expect higher *V\** values to result in greater fibrillar alignment, thereby increasing both *E* and *UTS*. For the consistent preparation of templated sheets, it was observed that increasing the oil droplet concentration beyond 2.3 v/v% in the WSS caused the sheets to tear when strained by the mandrel at *V\** = 10.

During the process of preparing ultrathin templated collagen sheets that includes microfluidic extrusion, washing, incubation, air-drying, and removal of oil droplets, the collagen concentration increased from an initial value of 5 mg/mL in the WSS by at least one order of magnitude (**Fig. 1C, Section S5**, Supporting Information).^[25]^ The volume concentration of droplets is expected to increase proportionally, assuming a uniform distribution of droplets, water removal by osmosis and drying, and mass conservation for collagen and oil (**Table S3**, Supporting Information).

The size distribution of the templating droplets was measured after passive production in a microfluidic flow-focusing device (model 3200152, Dolomite Microfluidics, Royston, UK). We considered two cases with different droplet diameters measured in the collection channel downstream of the site of droplet generation. In Case 1, the flowrates for the aqueous phase and oil phase were 30 µL h^-1^ and 480 µL h^-1^, respectively, which lead to an average droplet diameter of 5.0 ± 0.2 µm. In Case 2, the flowrates for the aqueous phase and oil phase were 15 µL h^-1^ and 720 µL h^-1^, respectively, resulting in a droplet diameter of 2.1 ± 0.6 µm. The droplet size distribution was wider in Case 2 than Case 1 because the target droplet size approached the lower limit of the droplet generator (1 ∼ 10 µm). The average diameter in Case 1 was selected to mimic the diameter of curvature of collagen fibers found in the artery wall.^[40]^ The average diameter in Case 2 was chosen to be comparable with the thickness of collagen sheets prepared under *V\** = 4.5 conditions (∼2 µm). Throughout the different processing stages of sheet fabrication, we observed broadening of the droplet size distribution, an effect attributed to droplet coalescence, particularly during initial mixing of oil-in-water emulsion with the collagen solution (**Fig. 2A**).

**Figure 2.**
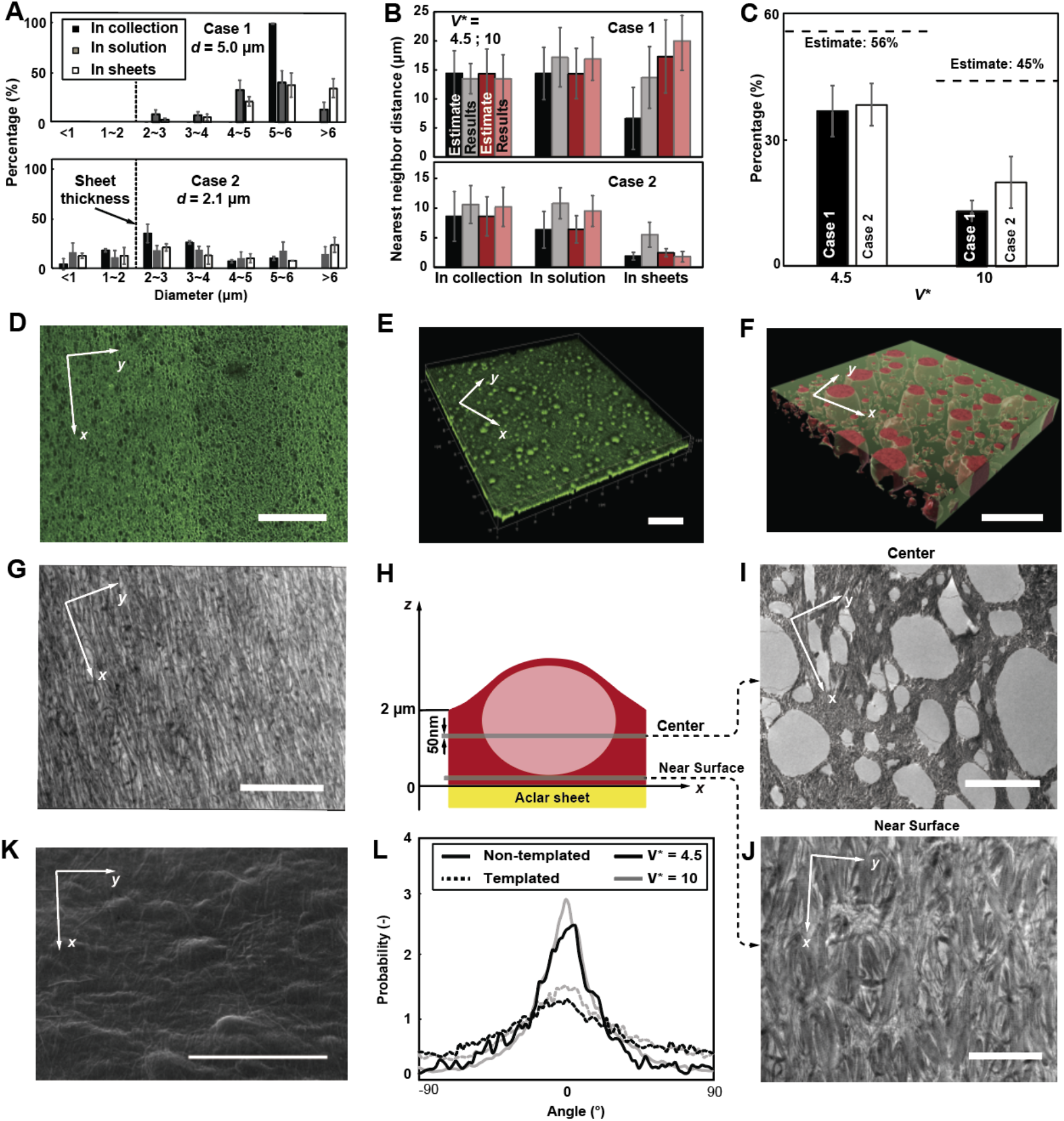
Morphological characterization of ultrathin templated collagen sheets. (A) Size distribution of templating droplets measured at different stages of sheet (*V\** = 4.5) formation and processing, for Cases 1 and 2. (N = 5) (B) Measured nearest-neighbor droplet distances (**Section S4**, Supporting Information). (C) Volumetric void fraction measured using confocal microscopy in templated sheets. (D) Stitched image from large-scale confocal scan of ultrathin, templated collagen sheet, *V\** = 4.5 (40ξ, scale bar: 50 µm). (E) Corresponding 3D rendered image (scale bar: 10 µm). (F) Local reconstruction of void distribution (red color) within ultrathin templated collagen sheets (scale bar: 5 µm). (G) Transmission electron microscopy (TEM) for x-y plane of non-templated collagen sheet control (*V\** = 4.5, scale bar: 10 µm). (H) Illustration indicating center and near wall planes where samples were prepared for characterization of the nanostructure of templated sheets. TEM images for planes near sheet (I) center and (J) surface (scale bars: 2 µm in I, 4 µm in J). (K) Scanning electron microscopy (SEM) image of templated sheet surface after post processing (*V\** = 4.5, scale bar: 5 µm). (L) Dispersion of alignment angles for templated ultrathin collagen sheets and control, evaluated from TEM images near surface.

The distributions of droplet size and nearest neighbor distance were analyzed from brightfield microscopy images. The results revealed an increasing mean droplet diameter and greater-than-expected nearest neighbor distance at different processing stages (**Fig. 2B**). We surmise that this is potentially due to droplet coalescence during the wet-spinning process, resulting in loss of droplets to the WSB bath, thereby reducing the droplet volume fraction (**Fig. 2C**). For the templated sheets prepared at higher *V\** values, a smaller droplet volume fraction was observed (**Section S5**, Supporting Information).

### 2.2. Morphological characterizations of templated sheets

Confocal microscopy in combination with electron microscopy (SEM and TEM) allowed us to investigate the ultrastructure of the templated collagen sheets. To investigate the void distribution within the templated collagen sheets, confocal microscopy (**Figs. 2D-F**) was performed using a 63x oil immersion lens (**Section S6**, Supporting Information). Z-stacks of the obtained images were reconstructed into 3D views. Image segmentation performed using a commercial microscopic image analysis software program (Imaris 10.0, Oxford Instruments/Bitplane, Schlieren, Switzerland) revealed the sheet internal void distribution at the microscale (**Fig. 2F**).

To better define the nanoscale organization of fibrillar collagen, transmission electron microscopy (TEM) images were obtained at the *x*-*y* plane, from 50 nm-thin precision cut slices (**Section S7, Figs. S3-S5**). Non-templated sheets served as a negative control and exhibited unidirectional alignment of collagen fibrils along the *x* direction, consistent with our previous report (**Fig. 2G**).^[25]^ Templated sheets were imaged at different *x*-*y* planes along the *z* direction (**Fig. 2H**). For specimen slices corresponding to *z* positions at the sheet centre, TEM images displayed the expected void regions (**Fig. 2I**). However, for slices near the sheet surface, no voids were observed (**Fig. 2J**). We speculate that the latter finding is due to the engulfment of templating droplets within the collagen sheet prior to evaporation, resulting in a continuous collagen layer at the surface. To support this hypothesis, we examined the surface morphology of the templated sheets after droplet removal using SEM, which confirmed our observations (**Fig. 2K**). The engulfment of collagen solution occurs during the early stages of sheet formation where the molecular crowding of collagen molecules formed a dense lamella that prevented the loss of droplets.

Directionality analysis (ImageJ, Release 1.45j, ImageJ, bundled with 64-bit Java8) was employed to quantify and visualize fibrillar alignment dispersion in TEM images (**Fig. 2L**). This allowed for the analysis of the orientation of structures by calculating the directional distribution based on pixel intensities. A distribution histogram was fitted to a Gaussian model, where peaks are identified as predominant directions. The resulting fibrillar dispersion spectrum was reported with respect to the x-direction.

According to the overall fibrillar orientation distribution, templated ultrathin collagen sheets displayed a broadened fibrillar dispersion, characterized by the full width at half maximum (FWHM) value of 64.7° for *V\** = 4.5 and 48.9° for *V\** = 10, as compared with 28.3° for *V\** = 4.5 and 22.5° for *V\** = 10 for non-templated sheets. The FWHM of a templated sheet produced at *V\** = 4.5 is approximately twice as large as that of a non-templated sheet, signifying greater dispersion induced by droplets at higher concentrations as compared to sheets produced at *V\** = 10. Additional characterization of fibrillar alignment is included in the supplementary information (**Section S8, Fig. S6-7**).

### 2.3. Biomechanical characterization of templated and non-templated sheets

Before biomechanical characterization, templated sheets were placed in a vacuum chamber connected to a vacuum pump via a cold trap at -80 °C to remove the templating oil (**Section S9, Fig.S8**). The complete removal of oil was confirmed using Fourier Transfer Infrared (FTIR) spectroscopy (**Figure S9**).

The tensile behavior of hydrated templated collagen sheets prepared in Case 2 was assessed and compared to that of non-templated sheets (**Table 1**) using a custom-built uniaxial tensile tester.^[25]^ Tensile tests were performed along both the *x-* and *y*-directions, where a least square curve of a continuous piecewise function, 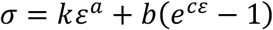, was fitted to the stress-strain datapoints.^[41]^ In the equation, *σ* represents stress, ε represents strain; *k* and *b* are fitting coefficients with units of Pascals; *a* and *c* are dimensionless coefficients (**Table S5**).

**Table 1.**
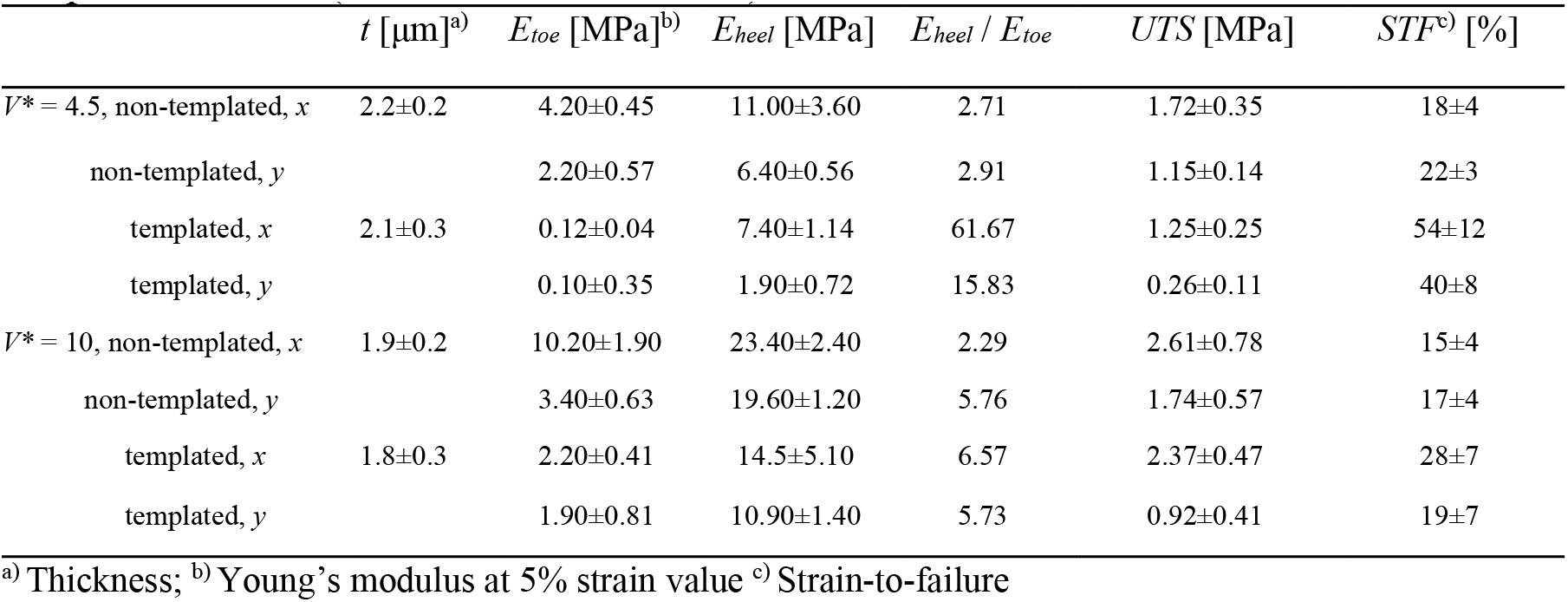
Summary of tensile properties of ultrathin, templated collagen sheets and nontemplated control. (N = 5 for all conditions)

To determine *E*_*toe*_ and *E*_*heel*_, the first-order derivative of the fitting function was evaluated at 10% and 90% STF, respectively.^[42]^ For each condition, stresses were measured for strain values that consecutively increased in 1/6 STF increments, up to the failure strain. **Figure 3** shows the average stress-strain curves for *V\** = 4.5 (**Fig. 3A**) and *V\** = 10 (**Fig. 3B**).

**Figure 3.**
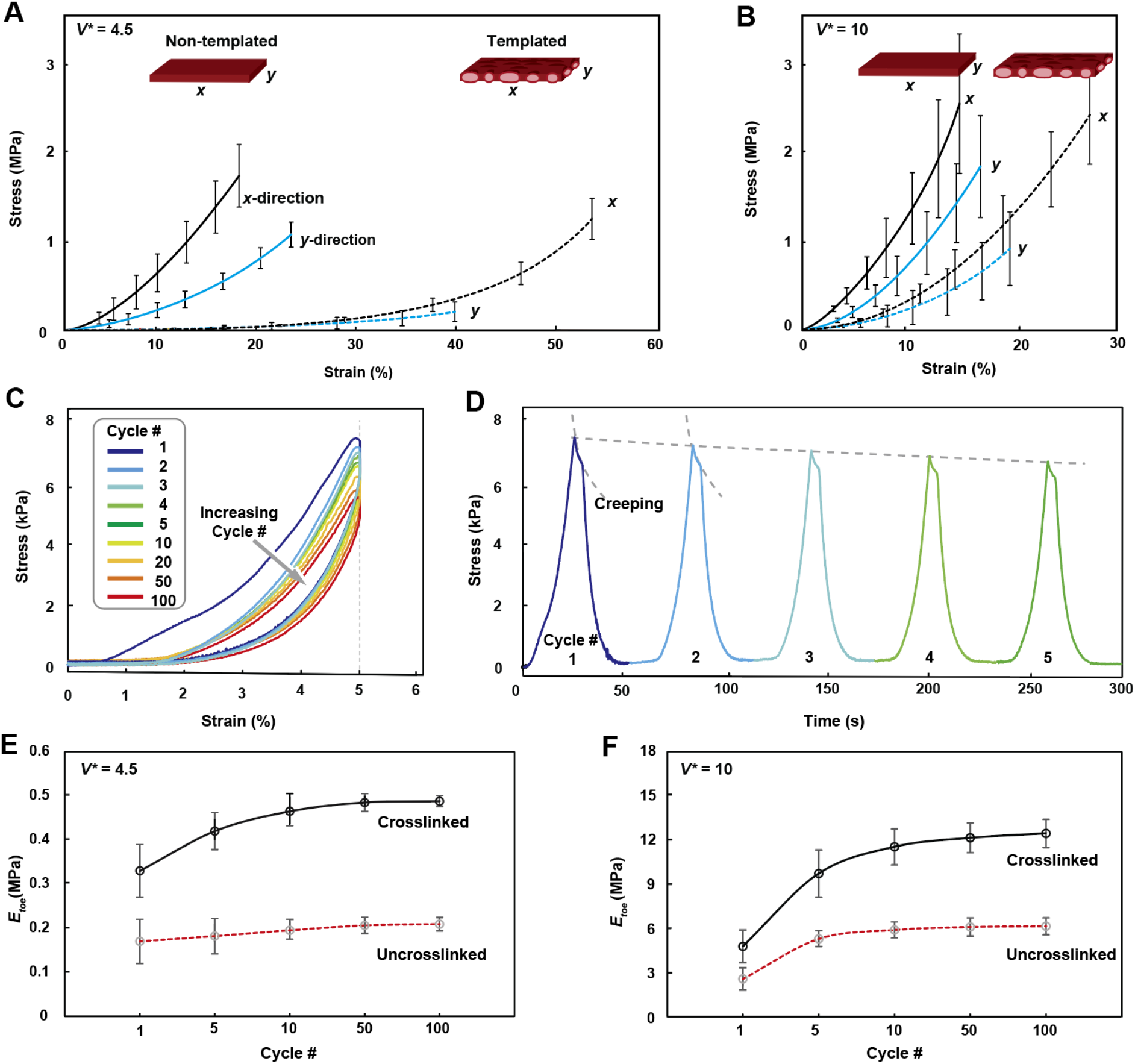
Biomechanical characterization of ultrathin templated collagen sheets. Tensile data obtained from uniaxial stretching experiments performed on templated ultrathin collagen sheets prepared at (A) *V\** = 4.5 (Case 2, N = 8) and (B) *V\** = 10 (Case 2, N=8). (C) Stress-strain measurements obtained for cyclic stretching of templated sheets prepared at *V\** = 4.5 up to 5% strain, 100 cycles. (D) Corresponding time-dependent stress measurements (moving average of 5 data points). Toe moduli for templated sheets prepared at *V\** = 4.5 (E) and *V\** = 10 (F) for non-crosslinked (gray) and 6 wt% genipin crosslinked (black) templated ultrathin collagen sheets (at 5% strain).

Both templated and non-templated collagen sheets exhibited anisotropic tensile properties. Under *V\** = 4.5 and *V\** = 10 conditions, templated sheets showed reduced thickness, *E*, and *UTS*, but increased *STF* as compared to non-templated sheets. Specifically, when produced at *V\** = 10, templating increased the STF in the *x*-direction by 60% while modestly decreasing *UTS* by < 10%. At *V\** = 4.5, with a higher void fraction, STF along the *x*-direction increased by > 100%, and UTS decreased by 27%. This indicates that a higher droplet concentration results in enhanced extensability of the templated sheets. At *V\** = 4.5, the ratio *E*_*heel*_/*E*_*toe*_ of templated sheets reached 62, favorably comparing to the behavior of arterial tissue.^[9]^

In accordance with studies on the tensile properties of cellular solids, the typical tensile stress-strain curves of polymeric cellular solids can be considered in three regimes, including a linearly elastic regime, where the cellular walls elastically bend or stretch, a stress plateau region, where cell walls plastically yield and pores collapse, and a densification ramp, where the pores are nearly fully collapsed, and the tensile load is borne by the cell edges and faces.^[36]^

In the case of ultrathin collagen sheets, the bending forces required to deform the edges of the materials are very small in terms of *E*_*toe*_. Consequently, the tensile curve of templated collagen sheets at low strain values does not exhibit a pronounced linearly elastic regime. Instead, the curve begins with the stress plateau region with low inclination (*E*_*toe*_), where internal pores collapse. As the applied strain increases and most pores become fully collapsed, the internal borders are increasingly aligned in the direction of stretch. This alignment leads to a marked increase in resistance to further elongation, resulting in a significant increase in Young’s modulus.

Tensile properties of the templated collagen sheets were evaluated in response to cyclic stretching for up to 100 cycles. During each cycle, the sheet was stretched to a 5% strain amplitude, and *E*_*toe*_ determined. For sheets prepared at *V\** 4.5 (Case 2), stress-strain (**Fig. 3C**) and stress-time (**Fig. 3D**) curves reveal hysteresis and a reduction in peak stress over time. During the holding period at the maximum amplitude, creep was observed, exhibited as a ∼15% decay in stress while maintaining 5% strain for 5 s. As the number of cycles increased, the magnitude of peak load reduction per cycle decreased. By the 100^th^ cycle, the peak stress decayed 25% as compared to the first cycle, which signifies the presence of irreversible deformation.

Strain-hardening was also observed (**Figs. 3E, F**). As compared to untreated templated collagen sheets, those crosslinked with 6 mg/mL genipin for 1 h at 37°C exhibit higher strength and a more pronounced strain-hardening effect. For templated sheets prepared at *V\** = 4.5, the increase in *E*_*toe*_ from the 1^st^ cycle to the 100^th^ cycle was 48% in crosslinked sheets, but only 23% in untreated sheets. For sheets prepared at *V\** = 10, which have a lower void volume fraction compared to those prepared at *V\** = 4.5, strain-hardening was more pronounced. The increase in *E*_*toe*_ from the 1^st^ cycle to the 100^th^ cycle as more than 240% and 300% for non-crosslinked and crosslinked sheets, respectively. These results suggest that under the same straining conditions, a greater portion of the collagen content underwent strain-induced plastic deformation. In contrast, sheets prepared at *V\** = 4.5, with a higher void ratio, are more capable of accommodating deformations at lower strains through fiber straightening. We postulate that strain hardening primarily occurs when fibrils are straightened and slide relative to each other.^[43]^ Tensile studies on multiple sheets stacked in parallel and at right-angles are included in **Section S10 (Fig.S10)**.

In summary, we successfully incorporated internal voids into ultrathin, aligned collagen sheets using templating oil droplets. The approach increased fibril dispersion as compared to non-templated collagen sheets (**Fig. 2L**). These nanoscale characteristics result in desirable non-linear tensile behavior mimicking native collagenous load-bearing soft tissues. At *V\** = 4.5 and Case 2 conditions (droplet diameter 2.1 ± 0.6 µm), we achieved *E*_*toe*_ of ∼100 kPa and *E*_*heel*_ of ∼1 MPa, with an *E*_*heel*_/*E*_*toe*_ of 62, consistent with the observed profile of cardiovascular tissue. We interpreted this non-linear tensile behavior in the context of a model of cellular solids, where *E*_*toe*_ and *E*_*heel*_ represent the stress plateau and densification regimes, respectively. In the stress plateau regime where voids collapse to mitigate stretch deformation, a higher void volume fraction allows for larger deformation without a significant increase in resistance to the tensile load. The more significant reduction in *E*_*toe*_ due to a higher void concentration results in a more profound *E*_*heel*_/*E*_*toe*_ ratio observed for sheets produced at *V\** = 4.5 as compared to *V\** = 10.

## 3. Conclusion

Templated ultrathin aligned collagen sheets were engineered with internal pores to mimic the strain-dependent tensile properties of native cardiovascular ECM. During templated sheet preparation, incorporating silicone oil droplets as a templating phase led to significant broadening of the fiber dispersion, achieving the desired J-shaped tensile behavior absent in non-templated collagen sheets containing highly aligned fibers. We confirmed that higher droplet volume fraction leads to higher fiber dispersion. In accordance with models of cellular solids, higher dispersion permits greater deformation without a substantial increase in the load in the stress plateau regime, contributing to a high *E*_*heel*_*/E*_*toe*_ ratio. At *V\** 4.5, pore templated sheets exhibited *E*_*toe*_ of ∼100 kPa and *E*_*heel*_ of ∼1 MPa, with an *E*_*heel*_*/E*_*toe*_ ratio of 62, similar to native cardiovascular tissues.

As compared to a zero-stress state, the circumferential pre-strain of arteries in response to physiologic blood pressure is ∼ 20%-50%.^[12]^ This stretch leads to a circumferential stress of ∼100-200 kPa.^[12]^ The tensile properties of templated sheets produced at *V\** 4.5 of 100 ∼ 200 kPa stress corresponds to 30% strain, which resides within a physiological pre-strain range.

Notably, cyclic stretching tests on templated sheets demonstrated a strain-hardening effect, particularly evident in crosslinked sheets where the amount of irreversible deformation increases as the number of cycles increases, quantified as the decay in peak stress at similar strain. This decay may potentially be attributed to lack of other ECM components that contribute to elastic tissue recoil, such as elastin and proteoglycans^[5, 44]^. These ECM components, especially elastin, dominates load bearing at small strain, ensuring resistance to plastic deformation.^[45]^ Future studies will expand this approach to additional ECM components, in order to more closely recapitulate the tensile behavior of cardiovascular tissues.

Overall, the described approach demonstrates significant potential for bottom-up biofabrication of load-bearing biomaterials to serve as soft tissue replacements. The ability to replicate the non-linear tensile behavior and maintain tensile strength that is physiologically relevant affords new avenues for the hierarchical design of biomaterials, particularly for applications in cardiovascular tissue engineering.

## 4. Material and Methods

### 4.1. Microfluidic preparation of templating emulsion

The templating emulsion was prepared using a commercially available microfluidic chip (model 3200152, Dolomite Microfluidics, Royston, UK). The detailed steps are described in (**Section S4**, Supporting Information).

### 4.2. Preparation of WSS and WSB

The WSB was prepared according to previously described.^[25]^ The WSS was prepared according to **Section S3**, Supporting Information.

### 4.3. Crosslinking of ultrathin templated collagen sheets

Genipin crosslinking solution (GCS) at 6 mg mL^-1^ were prepared by dissolving genipin in 1:1 v/v water and ethanol. Then, collagen sheets – after incubation in fiber incubation buffer – were immersed in GCS and placed in an incubator for 1 h.^[25]^

### 4.4. Morphological Characterization on ultrathin templated collagen sheets

The morphology of collagen sheets was imaged under wet condition were using a light sheet/confocal microscope (model Leica Stellaris 5), equipped with a water immersion lens (40× magnification, NA=1.1, article # 506307, WD = 0.65 mm) and an oil immersion lens (63× magnification, NA = 1.2, article # 506350, WD = 0.14 mm). The volume ratio of oil emulsion was analyzed using ImageJ software from the volumetric view constructed from z-stack scans. The surface morphology internal structure was studied with Hitachi SU1000 SEM and Hitachi HT7800 TEM respectively at the Nanoscale Biomedical Imaging Facility at The Hospital for Sick Children, Toronto, Canada. Detailed procedure of sample preparation is provided in (**Section S6**, Supporting Information)

## Contributions

Y.Z. contributed to the methodology, validation, formal analysis, investigation, resources, data curation, writing-original draft preparation and visualization. S.M. contributed to conceptualization, methodology, resources, writing - review and editing. B.W. contributed to software, formal analysis, investigation, and data curation.

E.L. contributed to methodology, resources, editing, and funding acquisition. A.G. contributed to conceptualization, methodology, resources, writing - review and editing, supervision, project administration, and funding acquisition.

## Acknowledgement

We thank Drs. Carolyn A. Haller and Richard Y. Cheng, for their suggestions. The cold trap was generously donated by late Prof. C.A. Ward. AG acknowledges funding from the Natural Sciences and Engineering Research Council of Canada (RGPIN-2017-06781) and the US (United States) National Institutes of Health (5-R01-EB032824-02). Device fabrication was performed at the CRAFT (Centre for Research and Applications in Fluidic Technologies) Device and Tissue Foundries are supported by the University of Toronto, the National Research Council of Canada, the Ontario-Québec Center for Organ-on-a-Chip Engineering and the Center for Advancing Neurotechnological Innovation to Application (Canada Foundation for Innovation, Ontario Research Fund).

Received: ((will be filled in by the editorial staff))

Revised: ((will be filled in by the editorial staff))

Published online: ((will be filled in by the editorial staff))

