## Supplementary Information for "Microfluidic Formation of Ultrathin, Handleable Collagen Sheets Exhibiting Toe-heel Tensile Behavior"

### 1. Literature Data for Elastic Moduli of Intact Tissues and Engineered Biomaterials

**Table S1. References for toe and heel elastic moduli of intact tissues and engineered biomaterials shown in Figure S1**

| Number in figure | Reference | $E_{toe}$ [MPa] | $E_{heel}$ [MPa] | *UTS* [MPa] |
|---|---|---|---|---|
| 1 | Daghrery *et al.*, 2023[1] | 0.05 | 0.4 | 0.07 |
| 2 | King *et al.*, 2022[2] | 0.2 | 1 | 1.6 |
| 3 | Rizzieri *et al.*, 2005[3] | 0.125 | 1 | 0.4 |
| 4 | Carotid artery[4] | 0.067 | 2 | 5.0 |
| 5 | Aorta[5] | 0.033 | 2.5 | 8.0 |
| 6 | Jugular vein[6] | 0.033 | 3.5 | 4.3 |
| 7 | Heart valve[7] | 0.18 | 3.1 | 2.9 |
| 8 | Bas *et al.*, 2017[8] | 0.1 | 7.0 | 0.6 |
| 9 | Sharabi *et al.*, 2021[9] | 0.5 | 2.5 | 4.2 |
| 10 | Kumar *et al.*, 2014[10] | 1.0 | 3.0 | 0.6 |
| 11 | Zhalmuratova *et al.*, 2019[11] | 1.0 | 4.0 | 9.0 |
| 12 | Zhang *et al.*, 2022[12] | 0.8 | 5.0 | 20 |
| 13 | Xu *et al.*, 2013[13] | 0.3 | 8.0 | 0.8 |
| 14 | Xu *et al.*, 2016[14] | 1.5 | 5.5 | 0.8 |
| 15 | Caves *et al.*, 2010[15] | 1.0 | 20 | 1.5 |
| 16 | Iliac[7] | 0.18 | 4.2 | 1.8 |
| 17 | Vena cava[7] | 0.21 | 9.6 | 1.7 |
| | This work, ultrathin templated collagen sheets | | | |
| 18 | $V^*$ = 4.5, templated, $x$ | 0.12 | 7.4 | 1.3 |
| 19 | $V^*$ = 4.5, templated, $y$ | 0.10 | 1.9 | 0.3 |
| 20 | $V^*$ = 4.5, non-templated, $x$ | 4.2 | 11.0 | 1.7 |
| 21 | $V^*$ = 4.5, non-templated, $y$ | 2.2 | 6.4 | 1.2 |
| 22 | $V^*$ = 10, templated, $x$ | 2.2 | 14.5 | 2.4 |
| 23 | $V^*$ = 10, templated, $y$ | 1.9 | 10.9 | 0.9 |
| 24 | $V^*$ = 10, non-templated, $x$ | 10.2 | 23.4 | 2.6 |
| 25 | $V^*$ = 10, non-templated, $y$ | 3.4 | 19.6 | 1.7 |

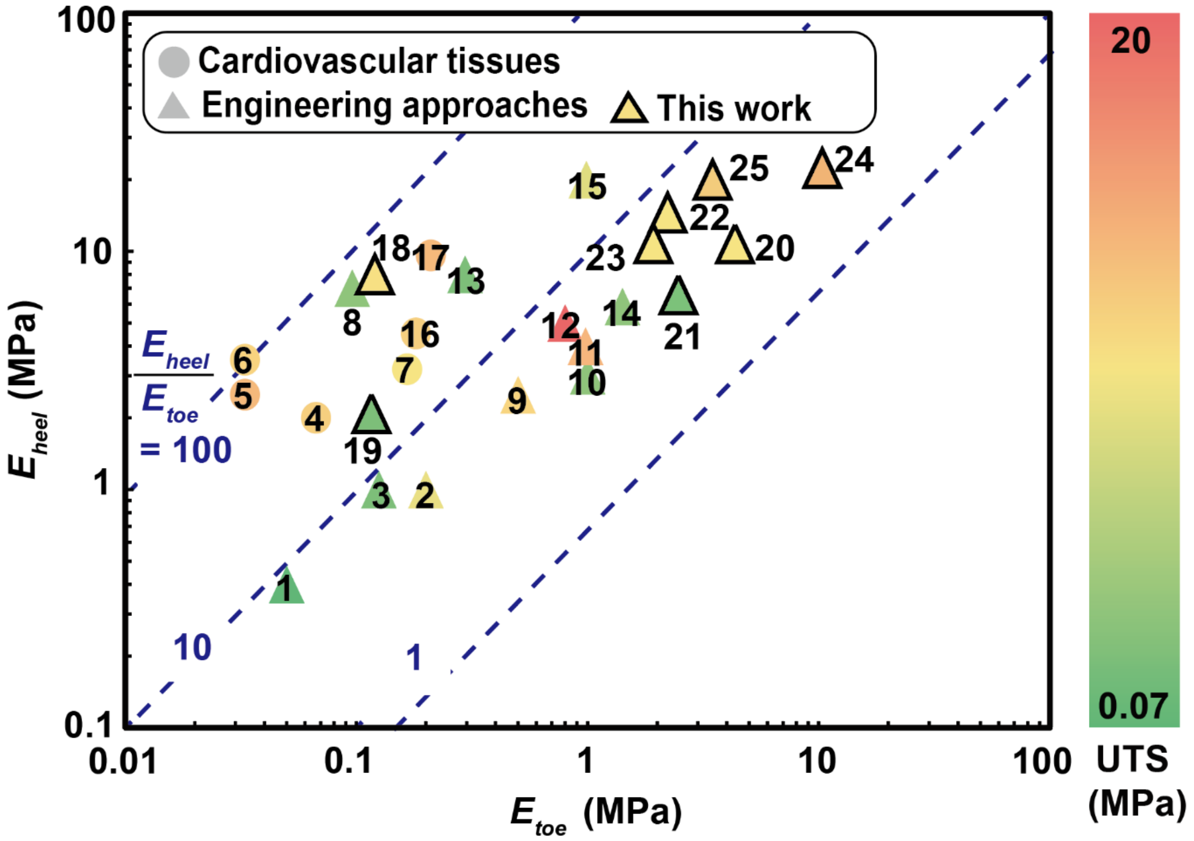


*Figure S1: Numbers in the scatter plot refer to references in Table S1.*

2. **Protocol Used for Extracting Rat Tail Type I Collagen**
    A) Spread rat tails (Cat# RT-T297, Rockland Antibodies & Assays, Philadelphia, PA, USA) on a paper towel for thawing during 30 min, until they assume room temperature, and cover with another towel to keep them clean and prevent dehydration of tendon during the thawing process.
    B) Adjust the pH of DI water to 2.0 using 37% hydrochloric acid (HCl) solution.
    C) Dissolve plier-pulled rat-tail tendons in HCl solution at room temperature overnight.[15] (dissolve the tendons at a concentration of 3 tails/L)
    D) Filter with 0.22 um filter using bottle filter setup (595 Bottle Top Filters – 500 ml capacity, MF75™ Series, Thermo Fisher Scientific, Waltham, MA, USA) at 4°C, change filter every 3 h.[16, 17] Store the filtered solution at 4°C.

The steps below are performed under sterile conditions

E) Salt out the collagen by adding 4.2 M sodium chloride solution to the filtered rat-tail solution at 1:5 volume ratio, keep stirring at room temperature for 15 min.
F) Centrifuge at 5000 rpm, for 4 h in 500 mL centrifuge bottles (Thermo Fisher Scientific Nalgene 3140-0500, Waltham, MA, USA), discard supernatant and dissolve the pellets in 500 mL HCl solution (pH = 2.0) at 4°C overnight.

G) Prepare 3 types of dialysis buffer solutions: (i). Phosphate buffer: 1.9 g sodium phosphate monobasic monohydrate and 9.34 g sodium phosphate dibasic anhydride in 4 L DI water; (ii). HCl buffer: pH 2.0 HCl solution; (iii). Pure DI water.

H) Transfer the collagen solution into an approximately 25 cm long, rinsed dialysis tube segment. Close the tube ends, submerge the tube in buffer. The volume of collagen solution in each 4 L dialysing buffer should be no more than 250 mL. Place a stirring bar at 10% of full speed and stir at 4°C for at least 4 h before changing buffer. The dialysis sequence is three times in buffer (i), three times in buffer (ii), and three times in buffer (iii).

I) After dialysis is complete, empty the tubes into 50 mL centrifuge tubes and freeze at -80°C overnight, tilt the tubes by approximately 30° to increase the surface area inside the tube. Maximum volume of solution contained in each centrifuge tube is 35 mL.

J) Lyophilize the frozen samples for 5 days or until ice pellets fully disappear at vacuum level 233 mTorr (30 Pa).

#### 3. Preparation of Wet Spinning Solution

To prepare the WSS in case 1, 50 mg lyophilized rat tail type I collagen was first added into 5 ml sterile pH = 2.0 water to yield 10 mg/ml collagen solution. Then, 3.6 ml 6.25 v/v% silicone oil emulsion (mean diameter 5.0 μm) and 1.4 ml pH = 2.00 water were added to the 10 mg/ml solution to yield the WSS (5 mg $ml^{-1}$ collagen and 2.25 v/v% droplets) for collagen sheet preparation. In case 2, after droplet generation, the 2 vol% emulsion was centrifuged in 15 ml centrifugal tubes (Falcon) for 30 min at 5000 rpm to separate the oil phase and water phase. Then, the aqueous phase was removed from the bottom of the tube using a 1 ml pipette to yield the same volume concentration at 6.25 v/v%, identical to that in case 1.

#### 4. Microfluidic Preparation of Templating Emulsion

The aqueous phase of was prepared from DI water and 3 v/v% polysorbate 80. The pH was then adjusted with 37 w/w% HCl to be 2.0. The oil phase was $5 \times 10^{-4}$ $m^2/s$ silicone oil (3176667, Sigma-Aldrich, Darmstadt, Germany). The two phases were filtered using 0.22 μm sterile filter before being filled into glass syringes (Hamilton™ HAM81420 glass syringe 10 mL, Hamilton, NV, USA). The glass syringes were then driven by PHD 4400 pumps (Harvard Biosciences) to pass the phases through a small quartz droplet chip with 10 μm channel depth and hydrophilic walls (model 3200152, Dolomite Microfluidics, Royston, UK). The flowrates for oil phase and water phase were 30 μL/h and 480 μL/h to yield 6.25 v/v% oil droplets with 5.0 μm mean diameter; 15 μL/h and 720 μL/h to yield 0.02 v/v% oil droplets with 2.1 μm mean diameter. The reagents were purchased from Millipore Sigma.

### 5. Increase in Collagen Weight Concentration and Oil Volume Fraction

For different values of $V^*$, the only difference is $v_m$, the linear velocity of the pulling mandrel. The relationship between $V^*$ and $v_m$ are related as

$$V^* = \frac{v_m - v_t}{v_t}, v_t = \frac{Q_B + Q_C}{WH} = 2.1,$$

where $Q_B = 400\ \mu l\ min^{-1}, Q_C = 4000\ \mu l\ min^{-1}, W = 35\ mm$ and $H = 1\ mm$.To estimate the collagen weight concentration in templated sheets, we need to first consider the collagen weight concentration in non-templated sheets. The calculation for sheets prepared at $V^*$ = 10 is provided as an example.

Consider 10 mL of collagen solution, given the initial collagen concentration used in wet spinning: 5 mg/mL, the total weight of dry collagen in the 10 mL solution was 50 mg. During extrusion where the flowrate of collagen solution was 0.4 mL/min, the time required to extrude 10 mL collagen solution was 24 min.

For a given mandrel pulling speed and corresponding $V^*$ value, together with the sheet thickness and width, the total volume (or length) of **hydrated** ultrathin collagen sheets can be estimated. Due to slippage between the emerging collagen sheet, the actual sheet length is lower than the estimated one. By comparing the actual length of the sheet produced, to the linear distance of the pulling mandrel travelled for, over 10 s, the slippage during sheet fabrication under $V^*$ = 4.5 and $V^*$ = 10 are measured to be approximately 24% and 33% respectively (Table S2). The yield can therefore be expressed to be "1-slippage%".

**Table S2. Slippage measurements during mandrel pulling mandrel pulling (N = 5).**

| *V** | *Linear distance travelled by mandrel in 10 s [cm]* | *Non-templated sheet length extruded in 10 s [cm]* | *Slippage* | *Templated sheet length extruded in 10 s [cm]* | *Slippage* |
|---|---|---|---|---|---|
| 4.5 | 12 | 9.1 ± 0.4 | 24% | 9.1 ± 0.4 | 24% |
| 10 | 23 | 15.4 ± 0.3 | 33% | 15.2 ± 0.3 | 33% |

For example, the total volume of non-templated, ultrathin collagen sheets produced from 10 mL collagen solution at $V^*$ = 10, assuming 33% slippage, can be calculated below given the sheet width of 12 mm and thickness of 1.9 µm:

$$\left(2.3\frac{cm}{s}\right) \times \left(24\ min \times 60\frac{s}{min}\right) \times (1.2\ cm) \times (1.9\ \times 10^{-4} cm) \times (1 - 33\%) = \ 0.51\ mL$$

The volumetric compaction from 10 mL collagen solution to 0.51 mL of sheet volume is:

$$\frac{10\ mL}{0.51\ mL} = 19.8 - fold$$

The average collagen weight concentration in sheets therefore is $\frac{50\ mg}{0.51 mL} = 99\ mg/mL$, and corresponds to a 20-fold increase in weight concentration compared to the concentration in the 5 mg/mL collagen solution.

Then, for templated sheets containing oil droplets, assuming that there is no droplet leaving the sheets, we expect the same fold change of oil volume concentration as the volumetric compaction. Given the initial volumetric concentration of oil droplets of 2.3%, the upper limit of the oil volume fraction is 2.3% ×20 = 46%. Based on sheet dimension and slippage measurements, the volume of templated sheets and non-template sheets produced from 10 mL of WSS is the same. However, in templated sheets, we estimate that 45% of the volume to be occupied by droplets. The volume occupied by collagen is therefore expected to possess a higher collagen weight fraction. In templated sheets prepared at $V^*$ = 10, the theoretical upper limit for the local collagen weight concentration is therefore $\frac{99\frac{mg}{mL}}{1-45\%} = 180\ mg/mL$.

The estimated fold change of the theoretical estimates of the local collagen weight concentration and the droplet volume fraction values under $V^* = 4.5$ and $V^* = 10$ are summarized in Table S2 below.

**Table S3. Estimated fold change in collagen concentration and void fraction calculation**

| $V^*$ | $v_m$ *[mm s$^{-1}$]* | *Thickness* [µm] | Yield [%] | Width [mm] | Volume [ml][a] | Collagen Weight Concentration in Non-Templated Sheets [mg mL$^{-1}$][b] | Fold Change | Estimated Void V%[c] | Collagen Weight Concentration in Templated Sheets [mg mL$^{-1}$][b] |
|---|---|---|---|---|---|---|---|---|---|
| 4.5 | 12 | 2.1 | 0.76 | 15 | 0.41 | 120 | 24 | 56 | 274 |
| 10 | 23 | 1.9 | 0.67 | 12 | 0.53 | 99 | 20 | 46 | 180 |

[a] Volume indicates estimate of total templated sheet volume attainable with 10 ml collagen solution during regular extrusion

[b] Collagen concentration in sheets after FIB incubation

[c] Theoretical void concentration induced by 2.3 v/v% oil droplets

For the distance calculation in sheets, since the droplet diameter is greater than the sheet thickness, the local structure can be illustrated in the drawing below, where the red square represents the sheet, the oil droplet is shown in yellow color. The intersection of the droplet and sheet volumes is shown in orange color.

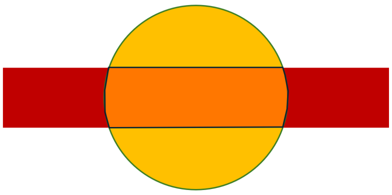

*Figure S2: Illustration of a droplet engulfed within a templated sheet segment*

In this illustration, the volume of the orange part can be calculated as the volume of the sphere deducted by the two spherical caps from the top and the bottom:

$$intersection\ volume = \frac{4}{3}\pi r^3 - \frac{2}{3}\pi\left(3r - \left(r - \frac{t}{2}\right)\right)\left(r - \frac{t}{2}\right)^2 = 37.18\ \mu m^3$$

Where $r$ denotes the droplet radius, $t$ is the thickness of the templated sheet without droplet. The ratio of the interception volume to the droplet volume is 56.8% for $V^*$ = 4.5 and Case 1.

Assuming volume conservation, estimates of the spatial nearest droplet-droplet distance can be obtained in the collection channel of the microfluidic device, in the WSS, and in templated sheets. Here, we estimate the distance by calculating the length of rectangular volumetric lattice which contains only 1 droplet assuming the droplet size and the spatial distribution of droplets are homogeneous. Sample calculations for $V^*$ = 10, Case 1 are shown below:

$$L_{in\ collection} = \sqrt{\frac{\left(\frac{unit\ volume}{number\ of\ droplets}\right)}{channel\ depth}} = \sqrt{\frac{\left(\frac{1\ cm^3}{\left(\frac{\frac{1}{17}cm^3}{\frac{4}{3}\pi(2.5\times10^{-4}cm)^3}\right)}\right)}{10\ \times 10^{-4}cm}} = 10.55\ \mu m$$

$$L_{in\ solution} = \sqrt[3]{\frac{unit\ volume}{number\ of\ droplet}} = \sqrt[3]{\frac{1\ cm^3}{\frac{0.023\ cm^3}{\frac{4}{3}\pi(2.5\times10^{-4}cm)^3}}} = 14.17\ \mu m$$

$$L_{in\ sheets} = \sqrt{\frac{\left(\frac{unit\ volume}{number\ of\ droplets}\right)}{sheet\ thickness}} = \sqrt{\frac{\left(\frac{1\ cm^3}{\left(\frac{0.45\ cm^3}{\frac{4}{3}\pi(2.5\times10^{-4}cm)^3}\right)}\right)}{1.9\ \times 10^{-4}cm}} = 17.06\ \mu m$$

The estimates of nearest droplet-droplet distance (L) for all conditions are summarized below:

**Table S4. Estimated nearest droplet-droplet distance**

| V* | | $L_{in\ collection}$ [μm) | $L_{in\ solution}$ [μm] | $L_{in\ sheets}$ [μm] |
|---|---|---|---|---|
| 4.5 | Case 1 | 10.6 | 14.2 | 6.0 |
| | Case 2 | 8.6 | 6.0 | 2.1 |
| 10 | Case 1 | 10.6 | 14.2 | 17.1 |
| | Case 2 | 8.6 | 6.0 | 2.4 |

### 6. Confocal Imaging

Prepared templated ultrathin collagen sheets were spread on a coverslip (140 μm thickness), incubated with 0.1mg $mL^{-1}$ fluorescein isothiocyanate-dextran solution and rinsed three times in DI water before imaging. An oil immersion lens (magnification 63, numerical aperture 1.2, corresponding depth of field 0.76 μm) was used on a scanning confocal microscope (model

Stellaris 5, Leica) was used to create the Z-stack (0.2 um per slice). All Imaging processes were performed using the 488 nm laser channel. After 3D reconstruction (Imaris 10.0, Oxford Instruments, Abingdon, UK), information that included the internal structural morphology, thickness, void concentration was obtained.

**7. Preparation of Templated Sheet Specimens for Transmission Electron Microscopy**

A) Spread hydrated collagen sheets on ACLAR® film (AGL 4458, Agar Scientific Ltd, United Kingdom), avoid wrinkles. Let air-dry inside a biosafety cabinet for at least 30 min.

B) Cut the ACLAR® film (AGL4458, Agar Scientific, Ltd, Essex CM24 8GF United Kingdom) with collagen sheets on top into rectangular pieces (maximum size 5 mm × 5 mm). Place each piece in one well of a 12-wellplate, add 1.5 ml fixation solution (2.5 v/v%) glutaraldehyde made in 100 mM cacodylate buffer (pH 7.2) per well and incubate overnight at 4°C.

C) Prepare osmium staining solution: 2 v/v% osmium tetroxide and 15 mg $mL^{-1}$ potassium ferrocyanide in cacodylate buffer (100 mM, pH 7.2).

D) Remove fixation solution and add the osmium solution and stain for 1 h, wash samples with double-distilled water ($ddH_2O$) for 5 min for 3 min each time at room temperature.

E) Prepare tannic acid staining solution consisting of 10 mg $mL^{-1}$ tannic acid in 100 mM cacodylate buffer (pH 7.2). Apply for 2 h, at 4°C, and replace with fresh 10 mg $mL^{-1}$ at 4 ºC.

F) Incubate in tannic acid staining solution for 2 h at 4ºC, wash sample according to step D).

G) Incubate samples with 2 v/v% osmium tetroxide in $ddH_2O$ for 40 min at room temperature and wash samples again thoroughly.

H) Incubate samples with 10 mg $mL^{-1}$ uranyl acetate (aqueous) at 4°C for 16 h, and then wash the sample according to step D).

I) Dehydrate specimen in graded ethanol: 30, 50, 70, 90, 100 v/v% ethanol in $ddH_2O$ for 10 min at each step. wash four times for 10 min each time at room temperature, and transfer samples to propylene oxide for 10 min at room temperature.

J) Place samples in mixtures of Agar100Hard and propylene oxide at room temp: first for 4 h in 30 v/v% Agar100Hard, then overnight in 50 v/v% Agar100Hard, and then for 1 h in each of 70, 90 and 100 v/v% Agar100Hard. Finally, repeat the 1-h treatment in 100% Agar100Hard twice more.

K) Cure samples in freshly prepared 100% Agar100 Hard in a labeled cylindrical aluminum mold for 72 h at 60°C, one specimen in each mold.

L) Remove block from the aluminum mold, shown below.

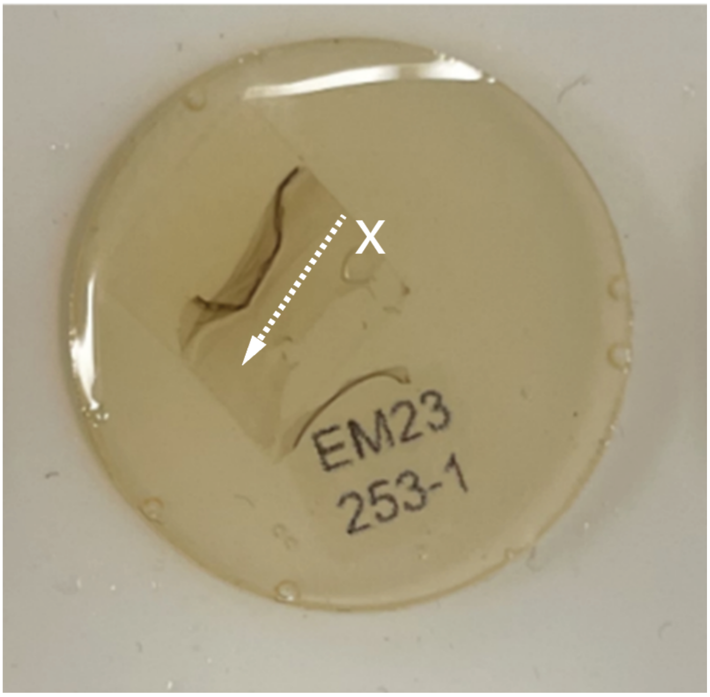


*Figure S3: Photo of an embedded templated sheet (V* = 4.5, case 2) spread onto ACLAR sheet.*

M) Isolate ~2 µm thick sections by manual cutting with a diamond trimming knife and place on a glass slide.
N) Stain the sections with 10 mg $mL^{-1}$ toluidine blue, rinse them in distilled water and dry the sections on a hot plate.
O) Examine the sections using a light microscope to verify specimen area, alignment orientation and section quality.
P) Prepare 50-nm-thin sections using an ultramicrotome, cut the section until the samples are captured.
Q) Collect the section on 200-mesh copper grids (Square Pattern 100 Mesh TEM Support Grids, AGG2100C, agar scientific, UK) and place under a TEM (Hitachi HT7800 RuliTEM, Hitachi High-Tech Canada, Canada).

Figure S2 shows an example of stitched TEM image ($V^*$ = 4.5, case 2), formed by capturing 25 images around the center. The images were then processed with ImageJ software to identify edges of the pores defined by the evaporated templating droplets (Figure S3). With the plugin, ilastik,[18] images were converted to binary format where enclosed shapes were identified as the result of oil templating, 255 in this particular case. The geometric center of each enclosed shape was identified using the particle analysis function in ImageJ. The nearest neighbor distances between droplets were then calculated using a MATLAB code shown below:

```
% Define the desired analysis image number here
file = 2;

% Code body
data = readtable("Centroid_00"+ num2str(file)+".txt");

X = table2array(data(:, 4));
Y = table2array(data(:, 5));

c2c_distance = [];
c2c_minimum_distance = [];

for i = 1:length(X)
    for j = 1:length(Y)
        x_dis = abs(X(j)-X(i));
        y_dis = abs(Y(j)-Y(i));
        D = ((x_dis^2) + (y_dis^2))^0.5;
        if D ~= 0
            c2c_distance = [c2c_distance, D];
        end
    end
    c2c_minimum_distance = [c2c_minimum_distance, min(c2c_distance)];
    c2c_distance = [];
end

c2c_minimum_distance = transpose(c2c_minimum_distance);
average_c2c_min = mean(c2c_minimum_distance);
min_c2c_min = min(c2c_minimum_distance);
max_c2c_min = max(c2c_minimum_distance);


Distribution_Results = array2table(c2c_minimum_distance);
Distribution_Results.Properties.VariableNames(1) = "Minimum Distance Distribution (um)";
filePath = "image_00" + num2str(file)+ "_Minimum Distance Distribution.csv";
writetable(Distribution_Results, filePath);
```

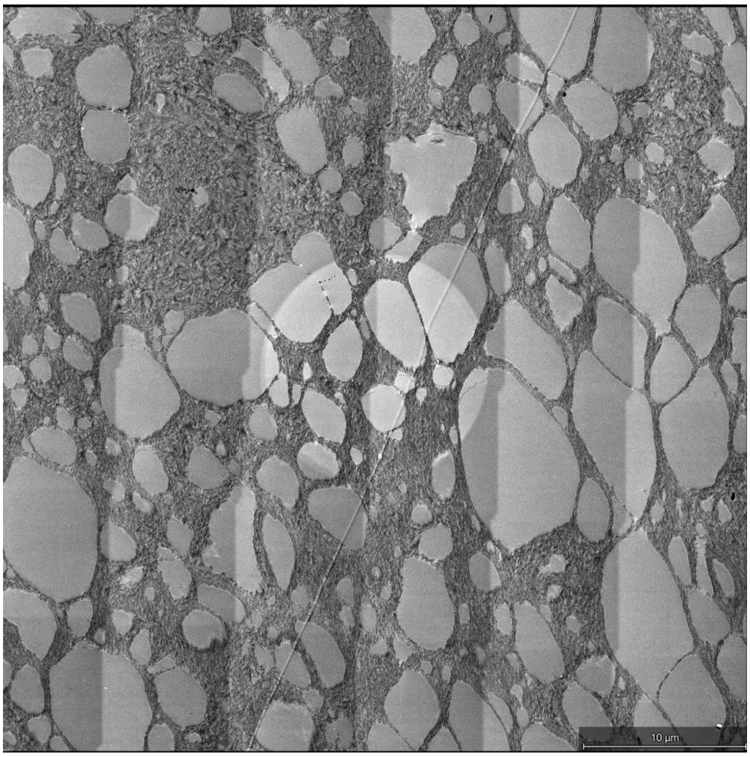


*Figure S4: stitched TEM image from 5×5 = 25 images, 5% overlap, V* = 4.5, case 2, xy-plane, case 2, 8-bit, width × height =21453× 21481 pixels, sample raw data.*

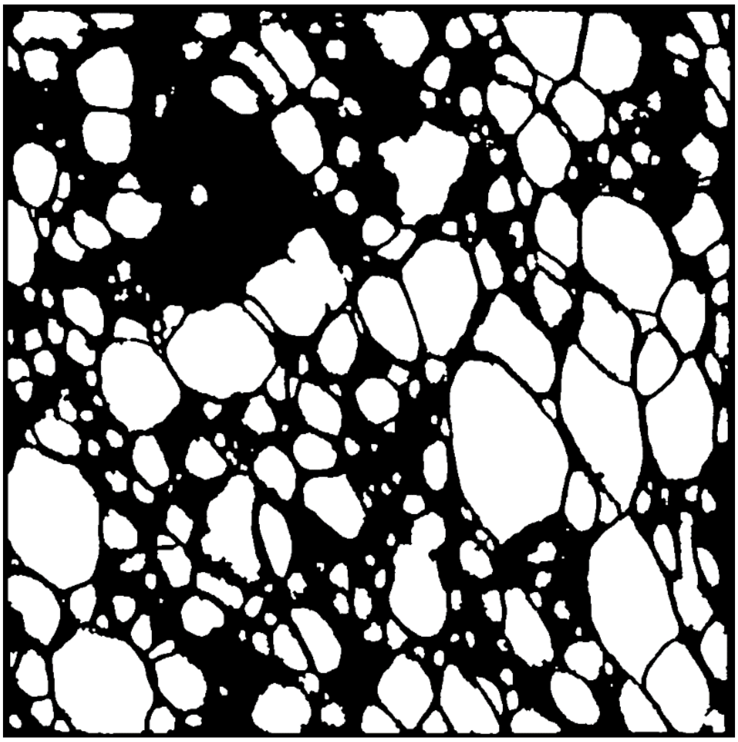

*Figure S5: Processed TEM image after thresholding and high-pass filtering (derived from Fig. S2)*

### 8. 2D Autocorrelation Analysis of TEM Images

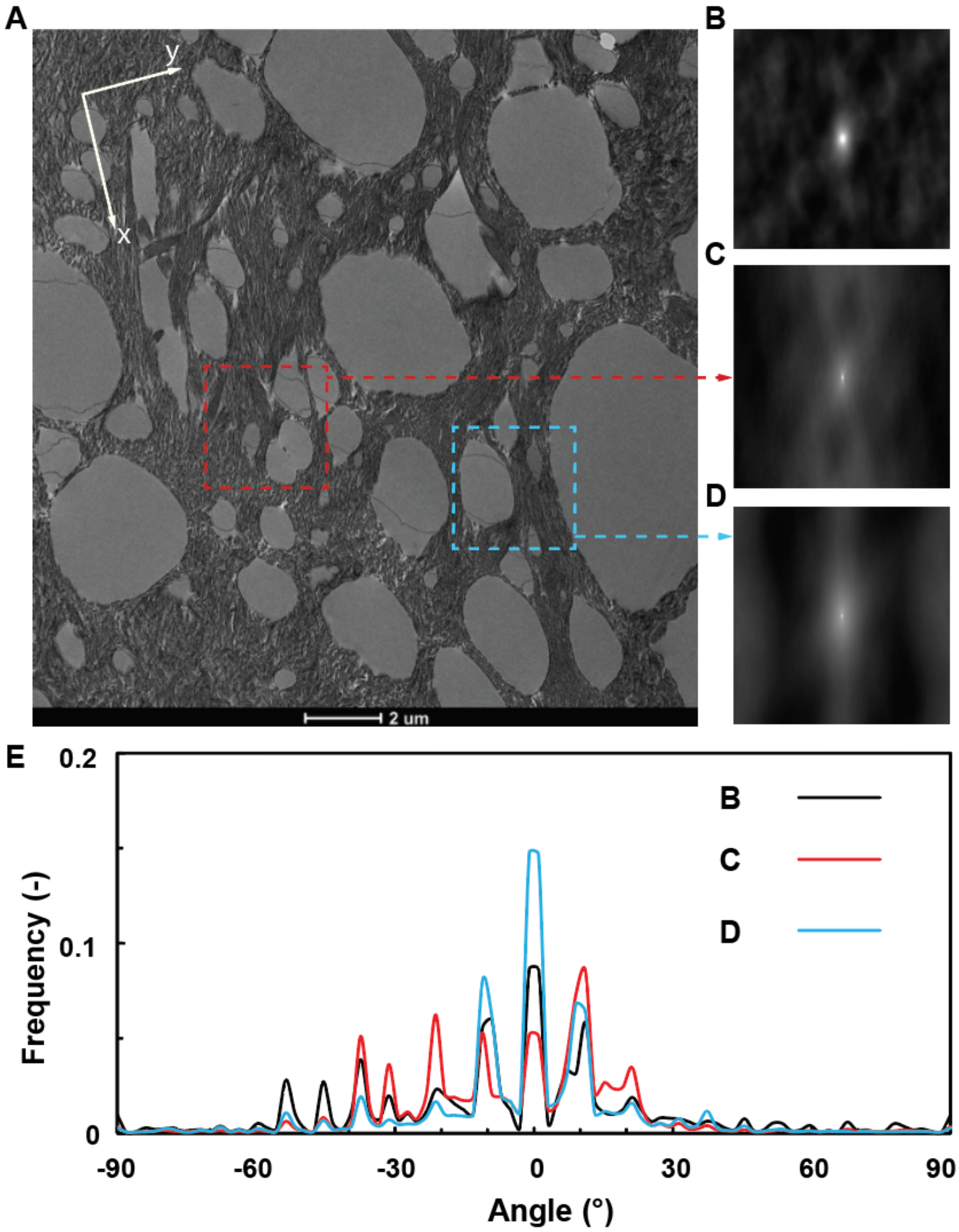


*Figure S6: A) Raw data of figure 2I (4096 × 4096 pixels). Colored squares refer to region used for local autocorrelation function (ACF) calculation. B) ACF of the entire image. C) ACF of the red square. D) ACF of the blue square. E) Alignment angle identification with respect to x-direction.*

2D autocorrelation analyses of the TEM images around the center along the z-direction of the templated sheets reveals the effect of templating on the dispersion of alignment angles (Figure S6). Comparing among the ACF's of the entire image and local images (Fig.S6B-D), the three images share most of the fibrillar alignment angles from -60° to 30° with respect to x-direction (-53°, -46°, -37°, -31°, -21°, -9°, 0°, 11°, 21°). Moreover, by comparing the peak values of curves C and D to B (Fig.S6E), we can identify the contributions of local fibrillar alignment to the entire image. To be detailed, the blue square region contributes to mostly directions along x-direction (peak values at -11°, 0°, and 9° are higher than the black curve); while the red square region contributes to angles including -46°, -31°, -21°, 9° and 21°. The other lower peaks in the black curve with higher peak height than the blue curve and the red curve indicates the fibrillar alignment angle contributed by regions other than the red and blue square enclosed regions.

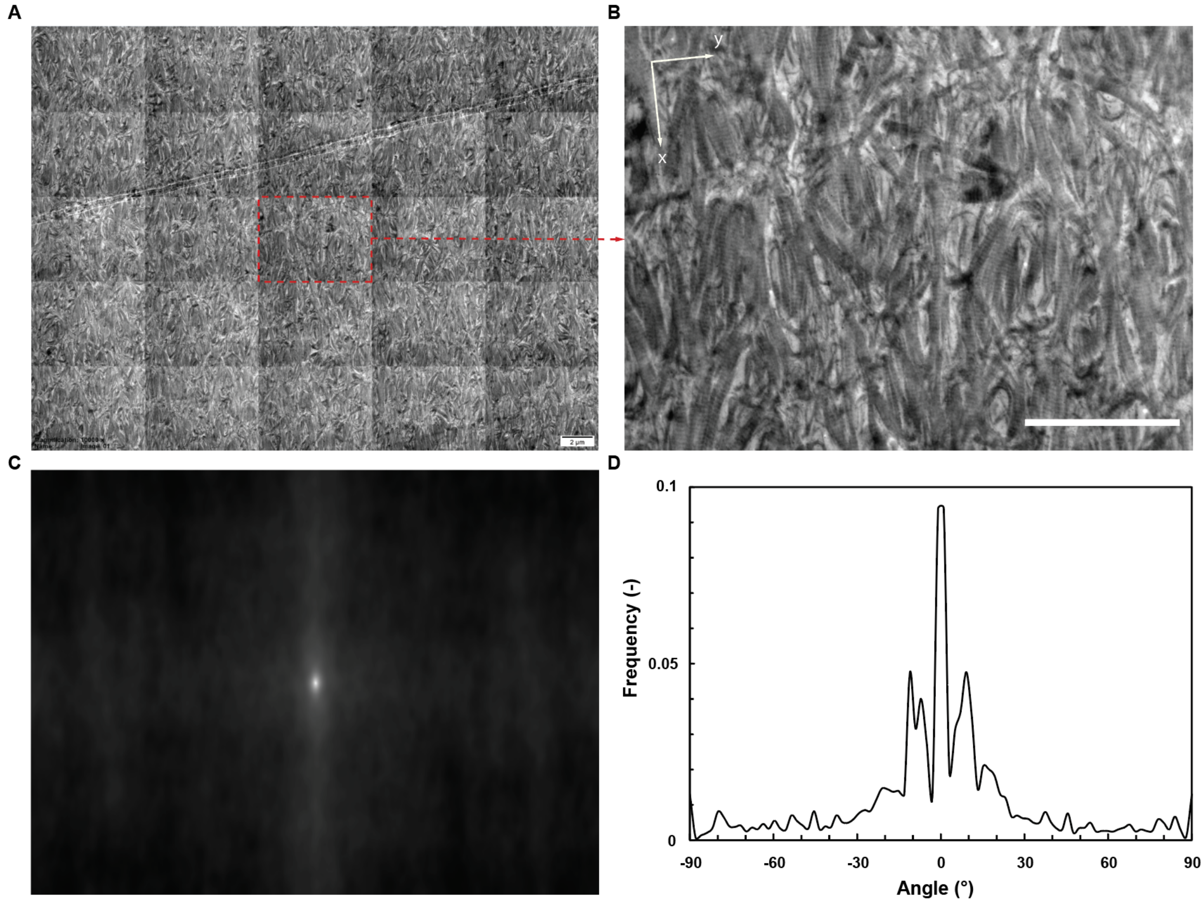


*Figure S7: A) Raw data of Fig. 2J (5 × 5 stitched images, 6000× magnification, 30kV beam, scale bar = 2 µm, 24-bit, 25307 × 18941 pixels, 72 dpi). The red square indicate region used for ACF in A) (scale bar: 2 µm). B) Zoomed-in view of the red square in A). C) ACF of B). D) Angle alignment with respect to the x-direction.*

The angle alignment patterns in TEM sample sections near the surface shows different characteristics due to the lack of voids (Fig. S7). The fibrillar alignment shows a broad peak between -11° and -3°, and 3° to 11° with respect to the x-direction (Fig.S7D). This reveals a higher degree of unidirectional alignment.

### 9. Evaporation of Template Oil Droplets

The oil removal was done using a vacuum setup previously reported by Singh *et al.*[19] Air-dried templated sheets were placed in a vacuum chamber connected to a vacuum pump (Edwards RV8, A654-01-903, Atlas Copco Group, Nacka Municipality, Sweden) via a cold trap (CentriVap -84 Cold Trap, 7460040, Labconco, MO, USA), vacuum tightened using clamp fitting (model 6653N15, McMaster-Carr, ON, CA). The absolute pressure in the chamber reached 0.05 mmHg (6.7 Pa) within 2 h. The vacuum was maintained 48 h for each oil removal session.

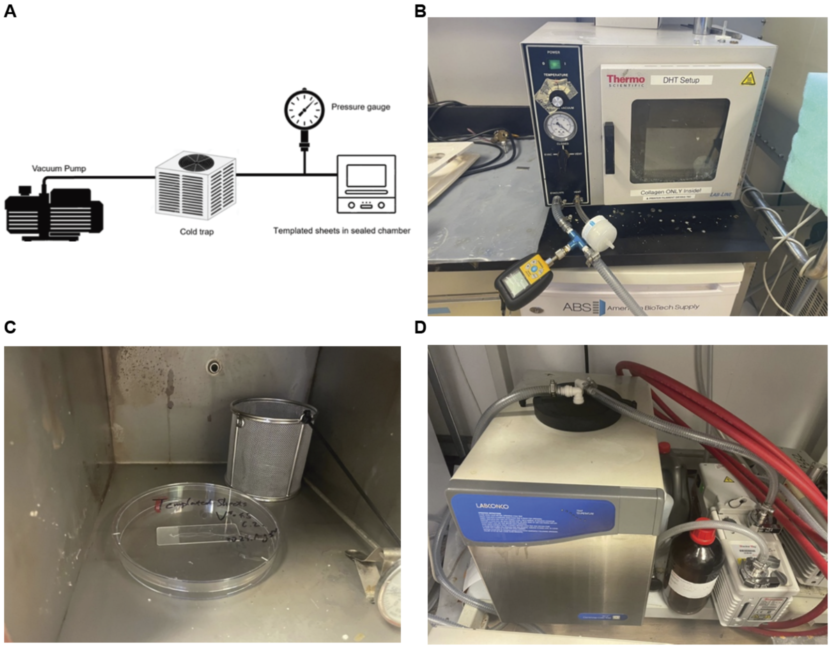


*Figure S8: Schematic (A) and photos of the oil removal setup B) Vacuum chamber; C) Sheet sample in the chamber; D) Cold trap and the vacuum pump.*

The templated sheets after performing oil removal were examined using attenuated total reflectance (ATR) FTIR (Nicolet™ iS50 FTIR Spectrometer, Thermo Scientific, Canada, penetration depth > 2.5 µm) and the peaks present in silicone oil were proved absent in the templated sheets, which confirms the complete removal of silicone oil using the method.

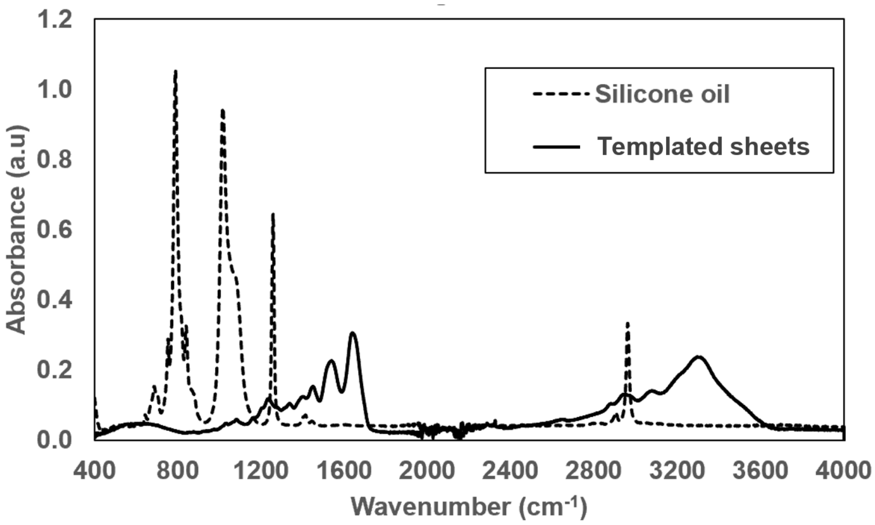


*Figure S9: FTIR scanning result indicating absence of oil in templated sheets (V* = 4.5, case 2) underwent oil removal.*

**Table S5.** Summary of fitting parameters calculated from the tensile data presented in Figure 3A, B, solving method GRG non-linear, initial input values: k = 5, a = 1, b = 300, c = 10.

| | $k$ [MPa] | $a$ | $b$ [Pa] | $c$ |
|---|---|---|---|---|
| $V^*$ = 4.5, non-templated, $x$ | 25.3 | 1.56 | 398 | 9.4 |
| non-templated, $y$ | 9.65 | 1.60 | 369 | 27 |
| templated, $x$ | 3.94 | 2.96 | 300 | 14 |
| templated, $y$ | 8.01 | 1.96 | 255 | 14 |
| $V^*$ = 10, non-templated, $x$ | 100.00 | 2.39 | 19.99 | 0.42 |
| non-templated, $y$ | 28.92 | 1.60 | 0.35 | 0.55 |
| templated, $x$ | 28.52 | 1.98 | 0.30 | 0.79 |
| templated, $y$ | 99.98 | 3.08 | 3.47 | 0.38 |

### 10. Tensile properties of double layer sheet stacks

Template sheets ($V^*$ = 4.5, case 2) were spread on top of each other at 0° and 90° with respect to the x-direction to form double-layer sheet stacks. The tensile properties of these constructs were measured by uniaxial tensile testing and compared to those of single layer templated sheets under the same extrusion conditions (Fig. S10).

When the two layers are at 90°, the difference between the stress-strain curves measured along $x$- and $y$-directions are insignificant. The increase in UTS of the 90° aligned double layers compared to single layer is insignificant but the STF decreases almost 40% from 54% to 33%. The Young's moduli including $E_{toe}$ and $E_{heel}$ both show increase, greater increase in the $E_{heel}$ leads to higher $E_{heel}/E_{toe}$ ratios over 90, which is 50% greater than that of single layer sheets (62).

When the two layers are stacked at 0° on top of each other, the increase in UTS compared to single layer sheets is insignificant in $x$-direction. However, the UTS along the $y$-direction increases over 80% from 0.2 MPa to almost 0.4 MPa. The strain to failure along $x$- and $y$-directions are 56% and 35% less. From the data regarding Young's moduli, the increase in $E_{toe}$ and $E_{heel}$ in $x$-direction are similar, which marks a similar $E_{heel}$ / $E_{toe}$ ratio (52) compared to single-layered sheets. In y-direction, a more pronounced increase in $E_{heel}$ makes the $E_{heel}$ / $E_{toe}$ ratio to be higher at 54, almost tripled from 19 – the ratio of single-layered sheets.

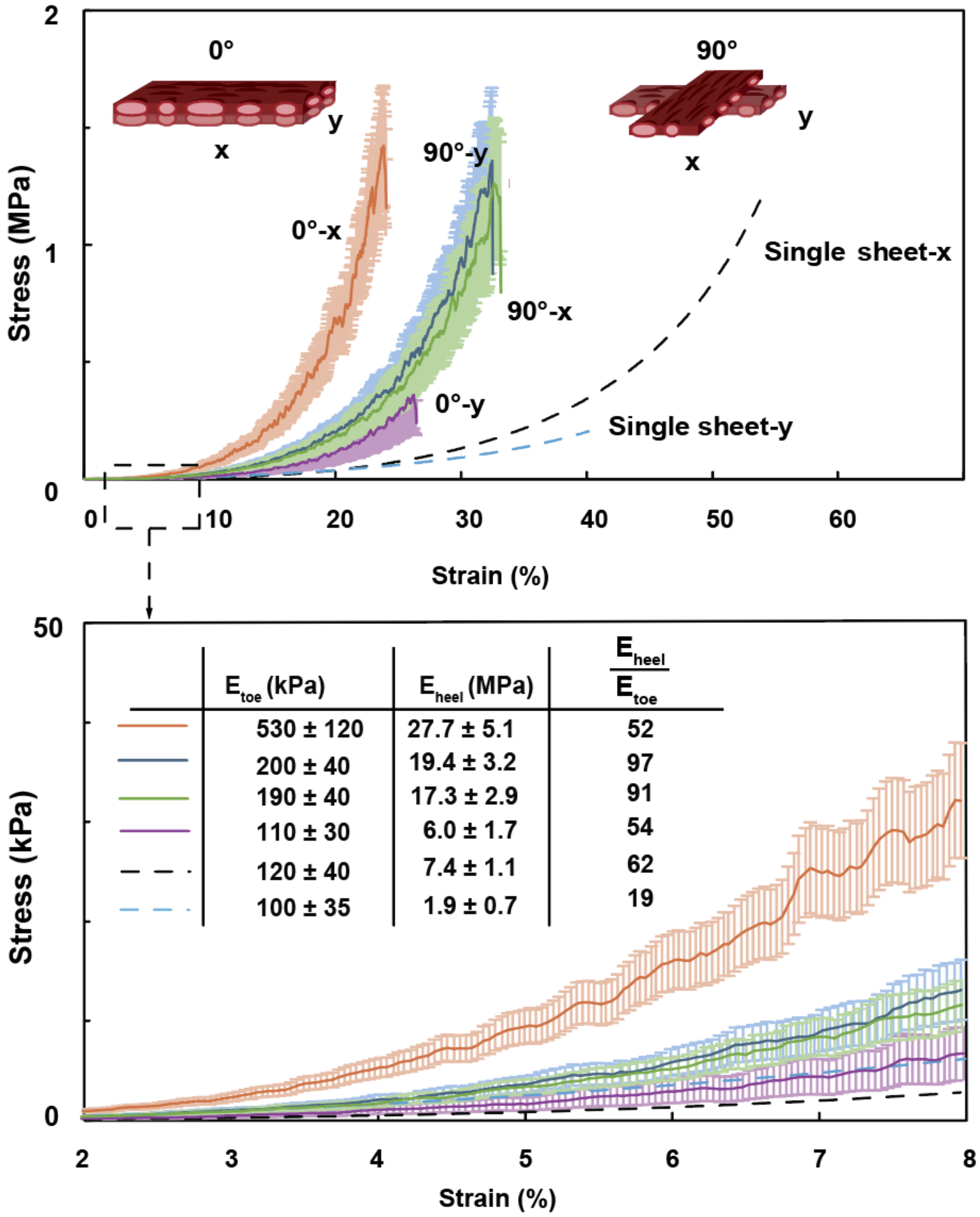


*Figure S10: Uniaxial tensile properties of double-layer sheet stacks (N=5) compared with single-layer sheets under V* = 4.5, case 2 conditions (data replotted from Fig. 3A).*
